# Parsers, Data Structures and Algorithms for Macromolecular Analysis Toolkit (MAT): Design and Implementation

**DOI:** 10.1101/605907

**Authors:** Gazal Kalyan, Vivek Junghare, S John S, Anupam Chattopadhyay, Pralay Mitra, Saugata Hazra

## Abstract

The structural information of biological macromolecules are stored in .pdb, .mm-cif and lately mmtf files and thus it requires accurate and efficient biological tools for various utilities. Here, we describe Macromolecular Analysis Toolkit (MAT) that parses .pdb, .mmcif and .mmtf files; and builds data structures from the input. This original program is written in C++ programming language to ensure efficiency and consistency to organize structural information in an integral way. The novelty of the program lies in the addition of new structure-based biological algorithms and applications. This package also stands out from other similar libraries by being 1) faster and 2) accurate. We also provide detailed comparison of available parsers on the whole PDB database. The parser of MAT is designed in such a way that it allows quick extraction and organized loading of the core data structure. The same data structure is extended to accommodate information from the .mmcif and .mmtf file parsers. Tokenization of the data allows the extraction of information from disordered text, making it compatible for accurate identification of the entities present in the .pdb file. Additionally, we add a new approach of performance optimization by creating a few derived data structures, namely kD-Tree, Octree and graphs, for certain applications that need spatial coordinate calculations. MAT provides advanced data structure which is time efficient and is designed to avail reusability and consistency in a systematic framework. MAT parser can be accessed online through bitbucket at https://bitbucket.org/gazalk/pdb_parser/.

## 1. Introduction

In the past decades, there has been a boom in the rate of growth of experimentally determined macromolecular structures. The three dimensional (3D) coordinates along with biological, molecular and experimental information of those experimentally solved biological macromolecules (protein, DNA, RNA) are stored in an online database known as Protein Data Bank (PDB) [1]. Each .pdb file is broadly divided into two parts: descriptive content and coordinate information. While coordinate information is well structured and easy to infer, most of the descriptive content is hidden as a running text. Looking into details, we find that every .pdb file of PDB is presented as an ASCII text file where each line consists of 80 columns and is terminated by an end-of-line indicator. In the header section, the information about protein, experimental details and additional structural attributes are mentioned. Every record starts with specific predefined keywords, however the information contained in them are predominantly textual in nature. Some records like ATOM, HETATM, MODEL, etc. are structured and follow strict rules facilitating easy parsing of its data. The file format itself is very primitive and with lot of limitations for automatic inferences. An attempt has been made to overcome some of the limitations in the .pdb file format by introducing the .mmcif format which is again very bulky and exhaustive. A recent advance has been made to introduce a binary file format as .mmtf [2] along with its parsers, which has also been integrated in this package. Our focus is to parse and load the .pdb, .mmcif or .mmtf file in our abstract data structure such that automatic information retrieval becomes effective and efficient.

A number of programs were developed as PDB Parsers and PDB based tools. Some early parsers are outdated due to either poor design or lack of maintenance such as BioC++[3], dsr-pdb[4] and PDBlib [5]. Python based programs like Biopython PDB Parser [6] does not provide the speed as needed for the present set of problems due to dynamic types and the use of an interpreter. With C++, we have much control on the code optimization to achieve high performance and speed. Many PDB parsers provide only basic data types and representations for a protein while others focus on specific applications. Victor C++ library [7] provides large-scale methods for structure manipulation which makes the code and data structure extra-detailed and heavy. ESBTL [8] is a library which allows handling of PDB data and also provides computational geometry methods. BALL [9] is another extensive library for algorithms. BiopLib is a C programming library for loading and manipulating macromolecular structures [10]. Finally, most of the parsers focus on the coordinate data explicitly and none describe handling of the header information especially REMARK section present in the .pdb file. However BiopLib [10] does provide substantial parsing of the header information which is generally neglected by other parsers. Haskell based library hPDB

The need of analysing protein structures i.e. their .pdb files is a high-end requirement of present era for making new drugs, analysing protein interactions, studying MD simulation intermediates and such other structure based studies. Now, .pdb files for macromolecular structures especially big ones are already disabled by RCSB and in this context it is very critical to have software platforms which could also read .mmcif and .mmtf files with good efficiency. Nevertheless, the existing structural techniques are still heavily based only on the .pdb file format.

## 2. Materials and Methods

### 2.1. Data set

For testing the features of our program, molecular structures for the test data set were selected from multiple sources:

1. The experimentally derived macromolecular structural data as available on RCSB-PDB was downloaded in the format .pdb and .mmCIF.
2. Intermediate MD simulation .pdb files.
3. Computationally modelled protein structures.
4. Macromolecular complexes resulting after molecular docking.

### 2.2. Tokenization of input file

Despite having format specifications for .pdb files, formatting deviations are often observed, as the .pdb file format is merely a plain ASCII .txt file. The lack of rigidity during creation and modification of .pdb files thus allows illegal token inclusions. This hinders the application of any automated software for data extraction based upon the guidelines specified for pdb format. Hence, the first step of our software is the tokenization. The process of tokenization is where the .pdb file data is broken into so called tokens or fragments. The lexical patterns in the file are identified using Flex [11] and the rule section of flex code is used to define the tokens. We utilize start conditions in Flex where we can conditionally switch between rules to tokenize the sections like REMARK, ATOM, HETATM, CONNECT, etc. In our program, we have two modes of tokenization: strict and casual. In casual mode, each record is read as one line per token and fed to next program for column-wise processing. It is difficult for computer programs to differentiate the identity of the atom without proper naming. Using strict mode in the MAT parser we have identified all the atoms that use non-standard atom names or disallowed characters in the file. Due to column wise distribution of the attributes in the .pdb file, there may be lack of delimiters and the values are compelled to merge. Hence, we use regular expressions instead of any delimiter for each column of the ATOM and HETATM record. As there are many sections in the .pdb files, we require various set of grammar rules for tokenization of each section which are elaborated in the Supplementary Tables S1 and S2.

The parser can be called in the following way, where the variable `optarg` is the file name for any of the .pdb/.mmcif/.mmtf file:

~~~
File::m_get_instance()->m_parse_file(optarg);
~~~

When you want to use the PDB ID of the structure and download it through the program, you can call the following function; here `optarg` being PDB ID:

~~~
File::m_get_instance()->m_download_pdb(optarg);
File::m_get_instance()->m_parse_file();
~~~

### 2.3. Syntax Tree formation

The individual tokens generated by the program are analysed and used by Bison (a parser generator) to identify the variables and add them to the internal data structure. To load data from PDB file in the data structure, we divide the file into various sections, start conditions from Flex program categorizes the section of the PDB file. PDB basically contain static representation of atoms in terms of X, Y and Z coordinates and header information related to that macro-molecular structure. A set of coordinates specifies the location of a point as an analogy for atom. However, we need to know the complete description of a single atom to describe its features that are mentioned in the .pdb file. The complete file is parsed by MAT PDB parser and the information (including atoms and header information) is stored in our data structure designed specifically for atoms. In our program, we avail the *Qt* library (https://www.qt.io/) for its container classes. The individual tokens generated by the program are analysed and used by Bison to identify the variables and add them to the internal data structure. Figure 1 explains the syntax tree formation and how components of the PDB are read and stored in our program.

**Fig 1:**
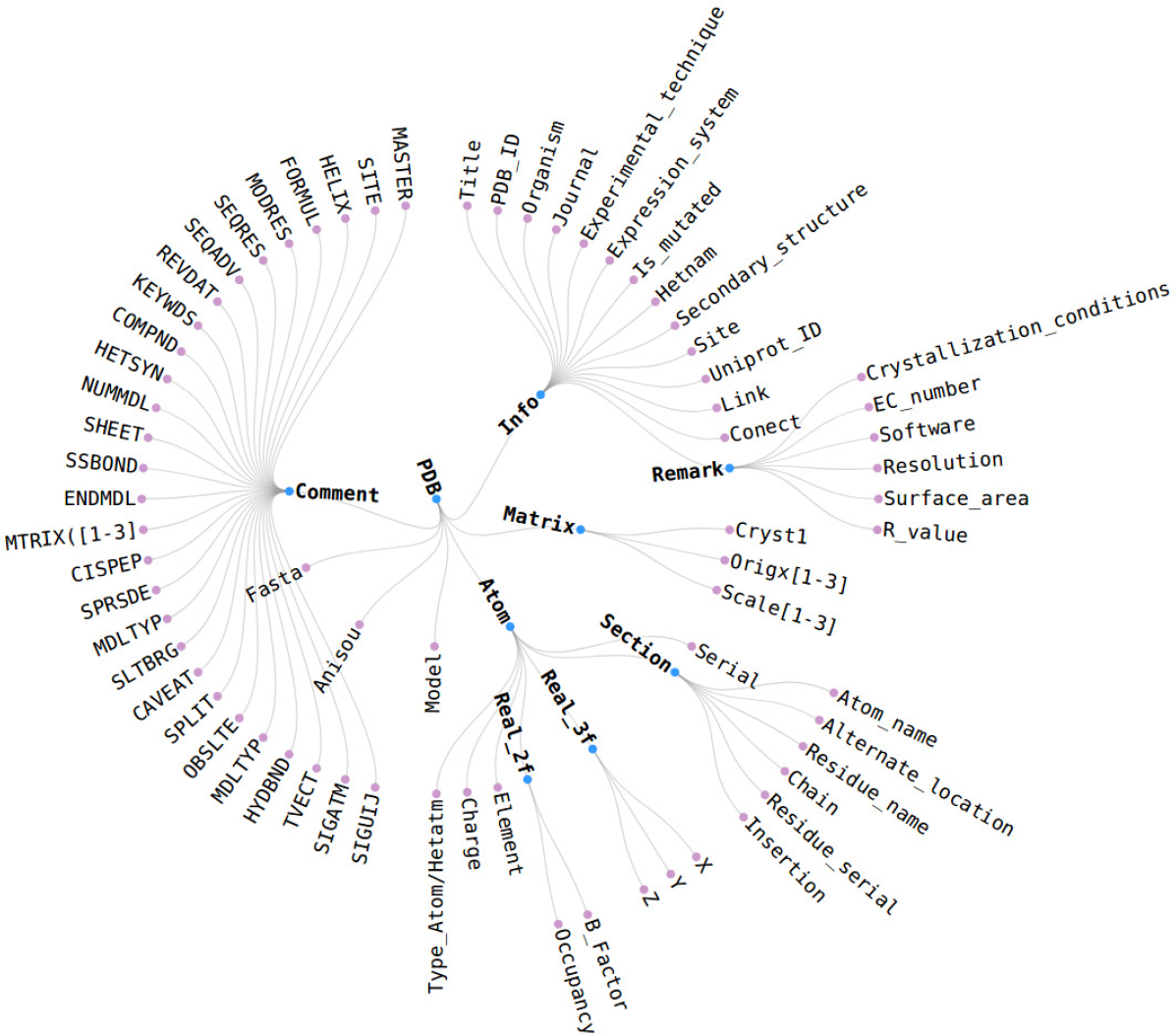
Syntax tree. This shows various components of the .pdb file. These are broken and identified into tokens and the macromolecular structure is maintained in our data structure for accessibility of atom/hetatm and accompanying information. The .pdb file attributes, info, remark, matrix, atom, model, fasta, anisou can be accessed using member functions provided. Some of data which is read but commented out is shown in comment section

### 2.4. Data structures

#### 2.4.1. Atom bucket

Atom bucket is constructed as an elementary abstract data structure using QList container. This data structure is simplistic in nature, all the atom(s)/ hetatm(s) are deposited in this primary bucket. The atom bucket class uses singleton design pattern which has only one instance. It has a global scope of accessibility. Atom is the smallest representation of protein structure in the PDB file; all the information and the properties that are associated to them are given for individual atom. Lastly, all the functions are which are available for a QList can also be used directly with the Atom bucket. On the whole, the atom bucket contains information on each of the protein points described in the .pdb file.

#### 2.4.2. Structural Hierarchy

For each level of structural hierarchy: atom/hetatm, residue and chain; we have a different bucket data structure which provides ease of accessibility to the information as they provide iterators for each level. Furthermore, every element of the level is represented as an object typed with its respective class. Since every level-element has been treated as a class, their properties, members and methods are also grouped separately. In doing so, we take in mind that all the groups at each level points to the same data. Therefore we can access, modify and use the information in atom-wise, residue-wise, chain-wise fashion. These designed buckets together make up the core data structure to handle the entire information of a protein system. Internally data structure looked like:

~~~
QList<Atom*>_m_Atom_Bucket;
QList<Residue*>_m_Residue_Bucket;
QList<Chain*>_m_Chain_Bucket;
~~~

And these are the iterator types to access these information

~~~
Atom_iterator
Residue_iterator
Chain_iterator
~~~

#### 2.4.3. Bonded Graph

A graph G(V,E) is defined to store the molecular structures, where V is the set of atoms and E is the set of covalent bonds among the corresponding atoms. The covalent bond information is stored as adjacency lists [12]. The interpretation of a chemical molecular structure is best represented by this data structure. It features identification of missing atoms in the structure and visualization of an atom in a structure even with multiple occupancies. For example, we can create a graph for all atoms in Atom Bucket as follows:

~~~
BondedGraph *m_Bonds
   = new BondedGraph((AtomRoot::m_get_instance())->m_get_atom_bucket());
~~~

### 2.5. Protein space partitioning

#### 2.5.1. KD-tree

k-dimensional (kD) tree is a binary tree with each node having maximum two children [13]. In case of protein structure, it is a 3D spatial tree which stores the 3D atom coordinates of a protein. It is used for searching the atomic nearest neighbors (NN) with respect to a query atom [14]. NN searches are performed very often while analyzing the structural aspects of the protein. kD-tree has been extensively used in related software such as ESBTL, BALL, Victor C++ library, etc.

We have iterated over the atom buckets to divide the protein space and store in kD tree. The root node represents the whole data space, at each level of the tree, the data is partitioned into two halves by the median point. This step is performed recursively for each dimensions X, Y and Z in the order of alternate turns. The median is selected from the list considering one dimension at a time by using quickselect algorithm which splits the dimension into two, for instance when a median is selected from X-axis the space is divided by the YZ plane. The left subtree would have values of the corresponding dimension greater than the median and the right subtree would have lesser values. This method creates a balanced tree which makes the search related to nearest neighbors fast. Figure 2 shows a simplistic example of a small kD-Tree. A few methods for NN search include k-nearest-neighbor, orthogonal search, circular search helping in the variety of biological problems. Algorithm 1 is for spherical search for kD-Tree radius search in atomic structures. It can be used and called in the steps mentioned below:

**Fig 2:**
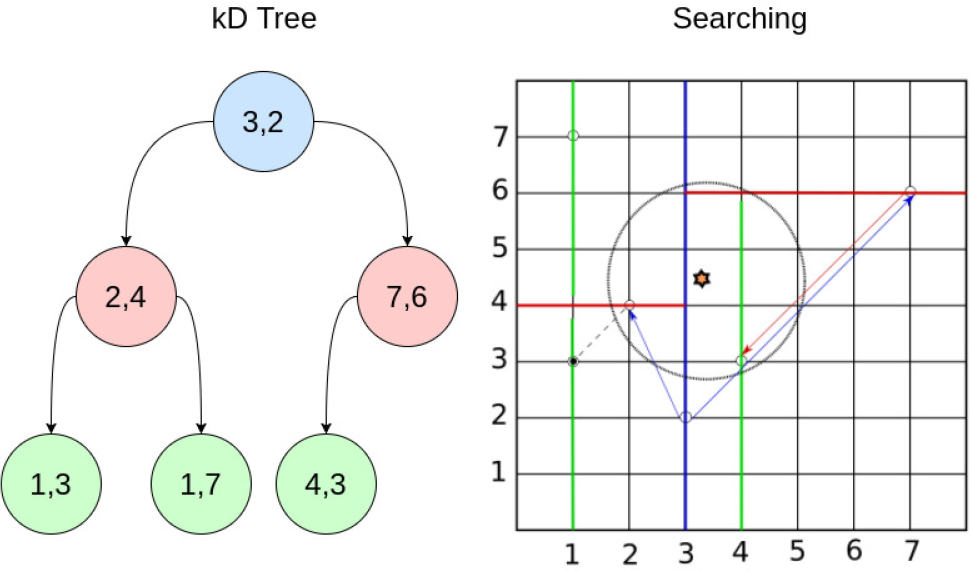
kD-Tree. The example of constructed kD-tree is shown with example of a 2-dimensional dataset. Searching for nearest neighbors of a query point(yellow star) is started at the root node of kD-Tree which is shown in blue. A recursive approach is then followed to check all the nodes and prune unnecessary sections.

~~~
KD_Tree* kd_tree1 = new KD_Tree(true);
~~~

Here, the variable `query_node` is the query point of type `Atom` and `range` is the radial range for the neighbour search to be given in Å.

~~~
Atom* query_node = new Atom(88457, “C”, “.”, “TYR”, “A”, 188, “.”,
  138.719, 33.258, 41.444, 1.00, 13.43, “C”);
kd_tree1->m_set_querynode(query_node);
QList<KD_Node*> KD_neighbors = kd_tree1->m_get_sphere_nn(query, range);
kd_tree1->m_print_list(KD_neighbors);
~~~

#### 2.5.2. Octree

Octree is a space partitioning tree with each node having exactly eight children. The root node contains the whole dataset and represents the orthogonal cuboid bounding box (voxel) for the data points as represented in Figure 3. These trees have been popularly used in the dynamic data sets and hence will be useful to deal with MD simulation data. The advantage of this tree is that insertion of the new data point does not need the tree rebalancing as each point finds its positions in this hierarchical spatial tree. For its construction, we first get the bounding box specifications for the data set i.e. the *min* and *max* values for X, Y and Z defining the enclosure for all the atom points. Then the space is divided recursively until it reaches the smallest node dimension possible i.e. voxel, as specified or until it contains single point in the cell. For *n* points, the insert function is called *n* times for insertion of each point in the octree data structure as explained in the Algorithm 2.

**Fig 3:**
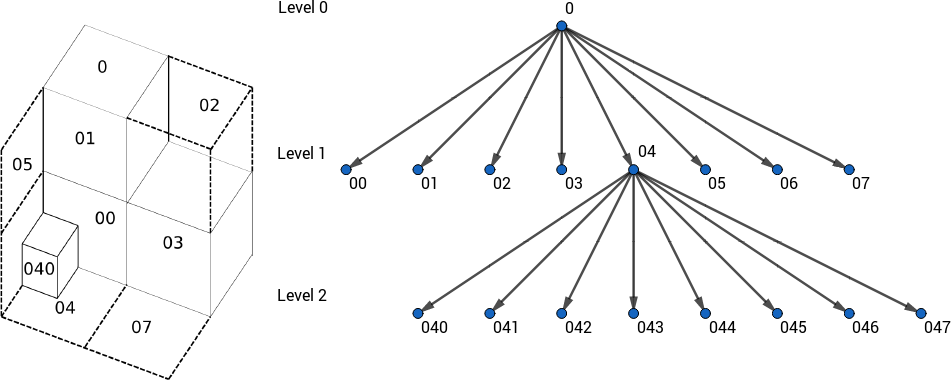
Octree nodes. The outer bounding box is set as the root node shown. Each spatial division splits the volume into its constituent eight separate volumes. 1^st^ division is shown in and in this level, we see the volume for 6th node (06) is not visible in the 3D cubic display. Each voxel is also uniquely represented and encoded as shown to identify its location.

We have implemented spherical search which is mostly used for molecular structures. The neighbor search is explained in Algorithm 3.

Octree contruction and Nearest-Neighbor searching can be used in following way. First we are giving an example of calling a function to ignore alternate locations of same atom:

~~~
Rule::m_get_instance()->m_keep_highest_occ();
Octree* octree1 = new Octree();
~~~

##### Algorithm 1: kD-Tree spherical search

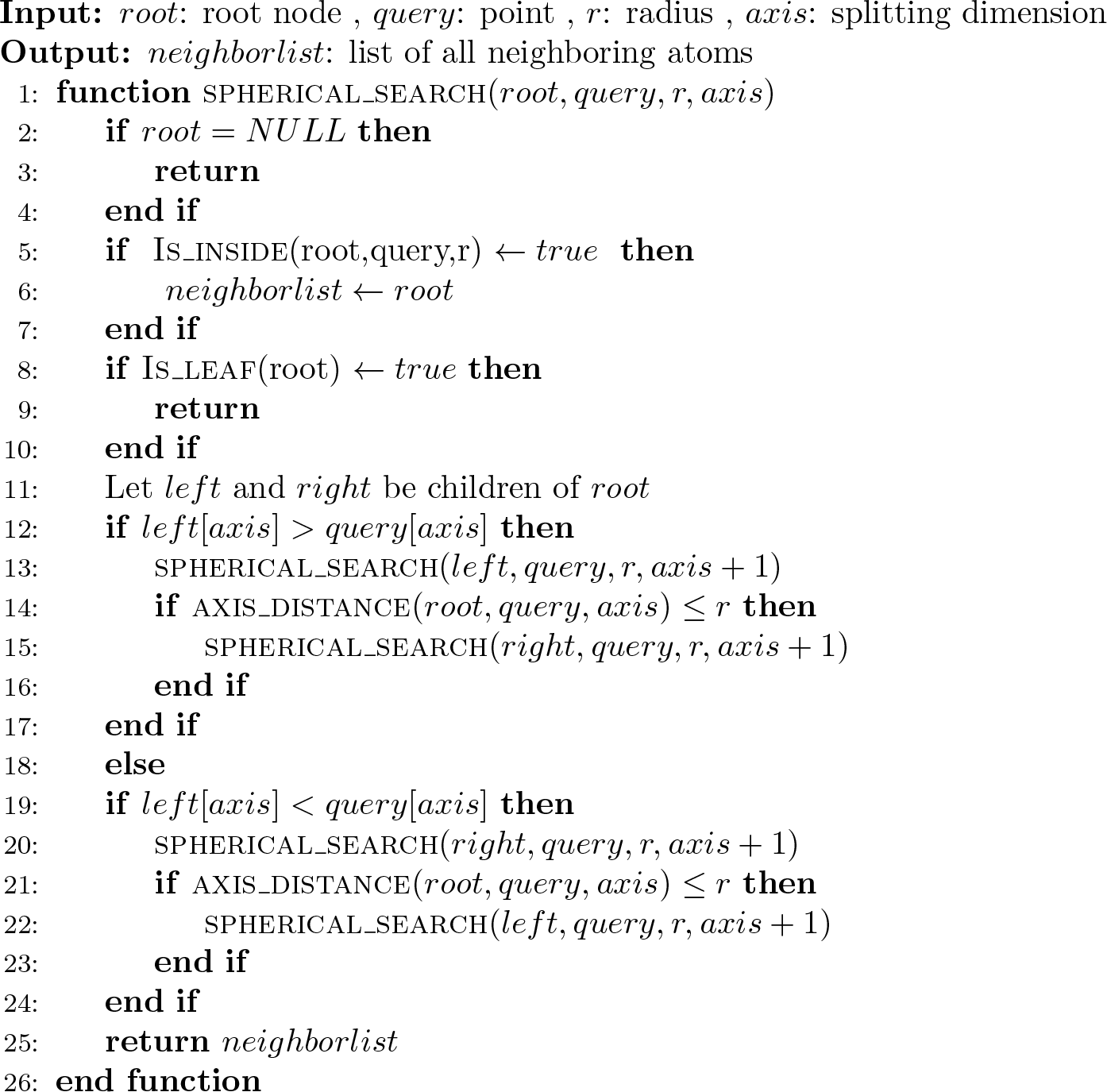

##### Algorithm 2: Octree construction

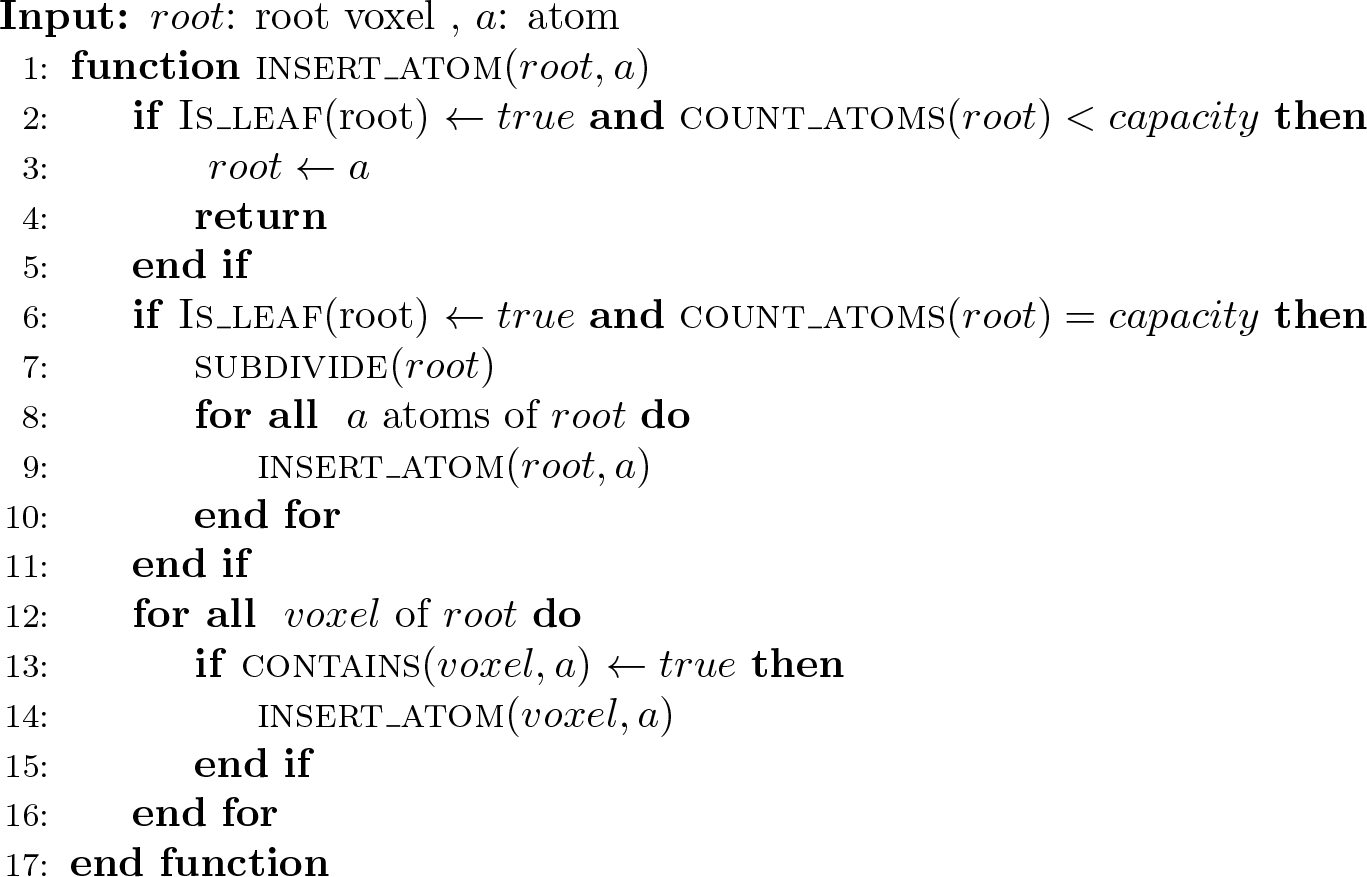

##### Algorithm 3

Octree spherical search

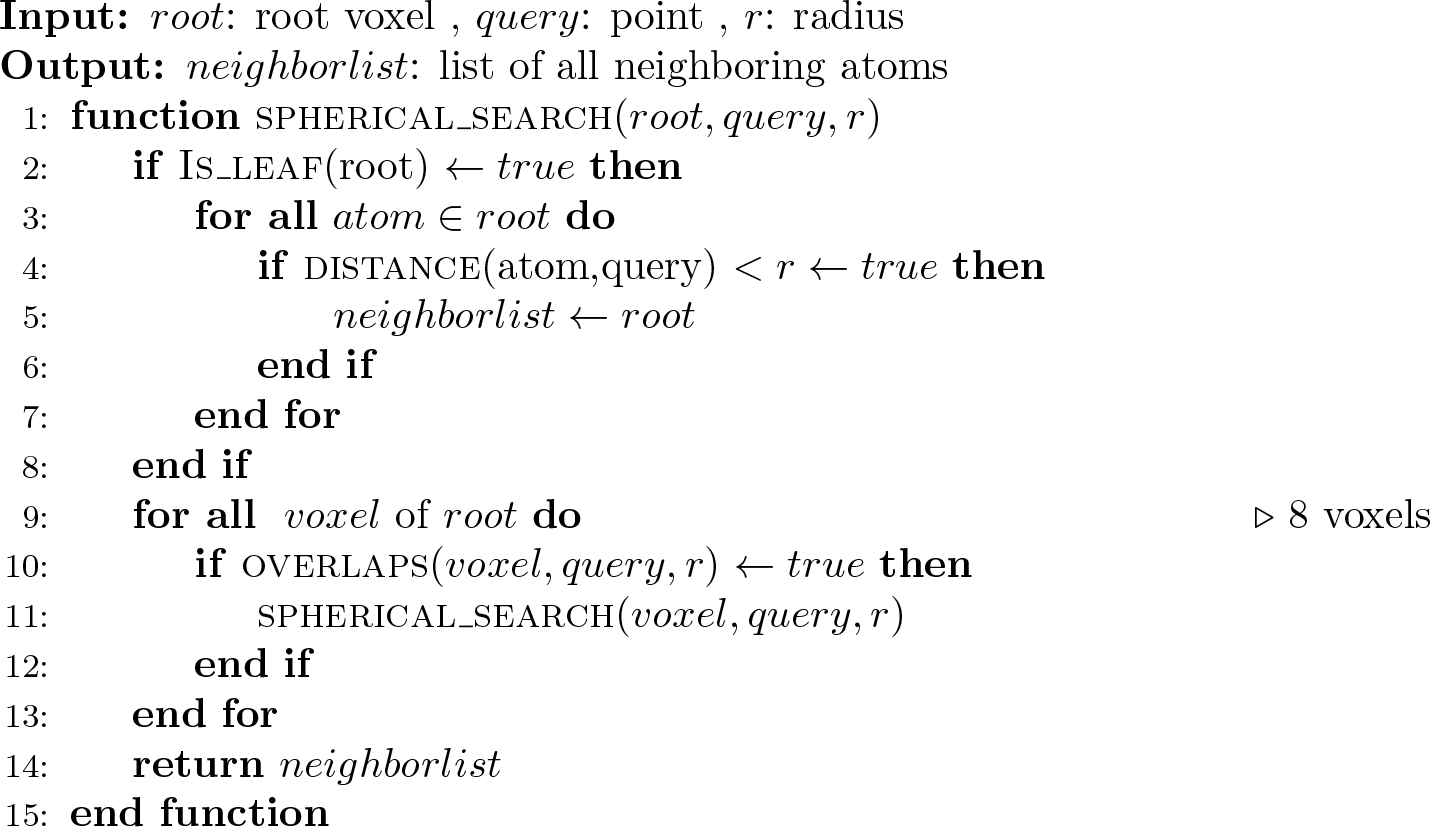

Here again, the variable the variable `query_node` is the query point of type `Atom` and `range` is the radial range for the neighbour search to be given in Å.

~~~
QList<Atom*> octree_neighbors =
  octree1->m_get_radius_neighbors(query, double(range));
octree1->m_print_list(octree_neighbors);
~~~

## 3. Results

### 3.1. Computational efficiency and speed comparison

For evaluating the performance and speed of our parser we have used the whole PDB database. Some files were incompatible due to bad formatting of the files (see section 3.2); however with regression testing and improvements in the code we are able to load all of the .pdb files successfully in our data structure with maximum accuracy. We also reported files which were non-standard with the pdb format. We have compared the runtime performance and efficiency of our software shown in Figure 4 with popular parsers. The benchmarks were made for MAT parser with programs: Pymol, BioJulia, BioPython, BioPerl, Victor C++, hPDB and ESBTL. It was observed to be fastest among all other packages that we tested for. The accuracy is based on the number of error occurrences while parsing the test data set as well as the ability to detect total number of atoms in the input file correctly plotted in the graph Figure 5. To compare the accuracy of count of atoms, the total number of atoms were calculated by counting the number of lines starting with ATOM and HETATM. This graph shows the number of files where the programs were either not able to read all the atoms in the file or they were not able to read the whole file.

**Fig 4:**
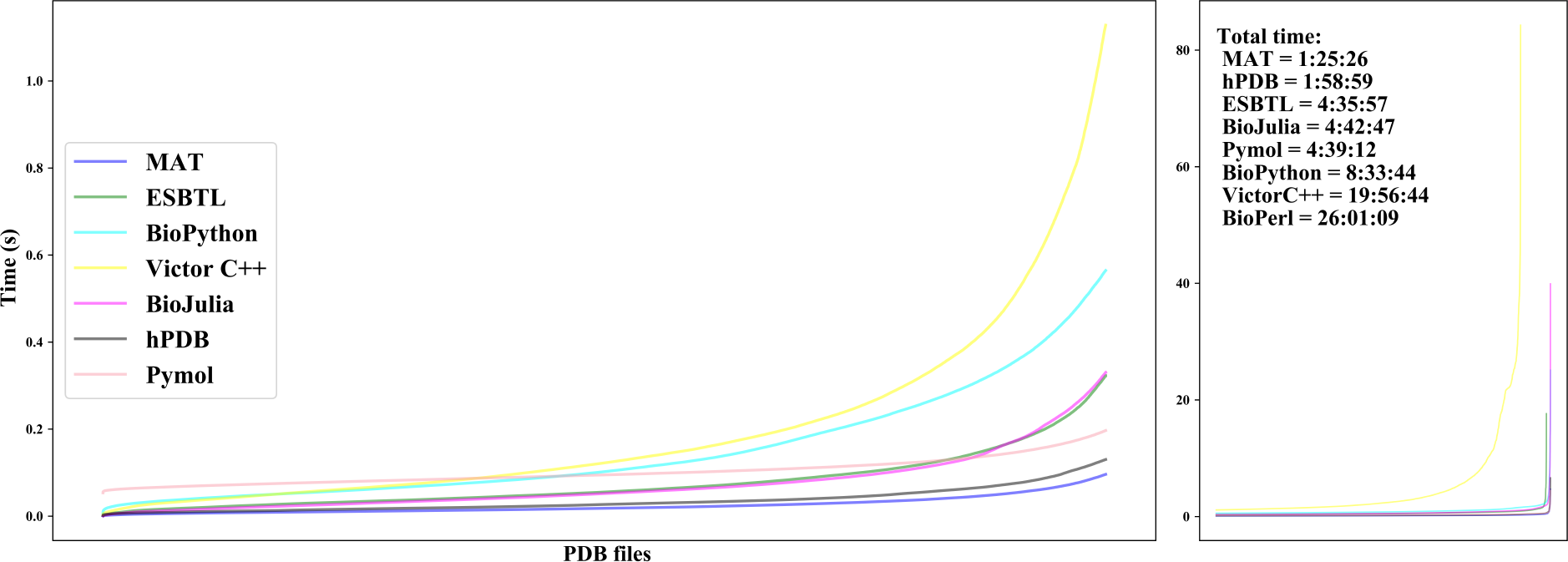
Speed comparison. The chart illustrates the speed of related software showing their relative performances on a UNIX based 64-bit system with Intel® Core™ i7-3770 CPU @ 3.40GHz × 8 with 8GB RAM. This test was made on 145787 number of file and the values were sorted based on ascending time. BioPerl has been skipped to obtain a clean graph. Total time taken for parsing the whole database is given in the format HH:MM:SS.

**Fig 5:**
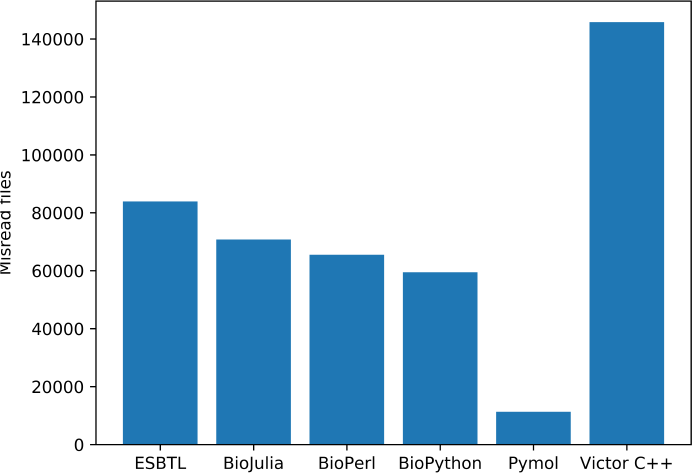
Parsing total atoms in a file. This graph shows the number of instances of misread files. MAT Parser and hPDB were completely accurate and did not show any errors. Although rest of the parsers encountered misreading of files as shown with BioJulia, ESBTL and Victor C++ throwing errors on 2, 132, 964 files respectively..

The algorithms of nearest neighbor search for our data structures were compared and tested for their accuracy and speed using Google Test [15]. It was found that both the kD-tree and Octree are fairly faster than simple linear search. The neighbors were exactly accurate in all the test cases. Furthermore, Figure 6 shows speed comparison of different voxel capacities of Octree and kD-Tree, the latter being fastest. We found that voxel size of 3 was optimum and if we start to increase the voxel capacity more than 32, we see a noticeable difference in the speed. We have used kD-Tree extensively in finding networks of non-covalent interactions such as salt bridge and hydrogen bonds. It is advantageous to use this data structure especially where there is a requirement of frequent neighborhood search queries multiple times, for example, interaction studies. Secondly, it is stated that kD-Tree can only be used with static data set. If we consider semi-dynamic or dynamic data sets, it is advantageous to use Octree. Therefore for molecular dynamics and energy minimization, Octree can be used. It is so because the updation in this data structure does not involve altering the whole data set but it only refer to positional changes inside the local reference frame of the bounding box. The benchmarks are given in the supplementary material.

**Fig 6:**
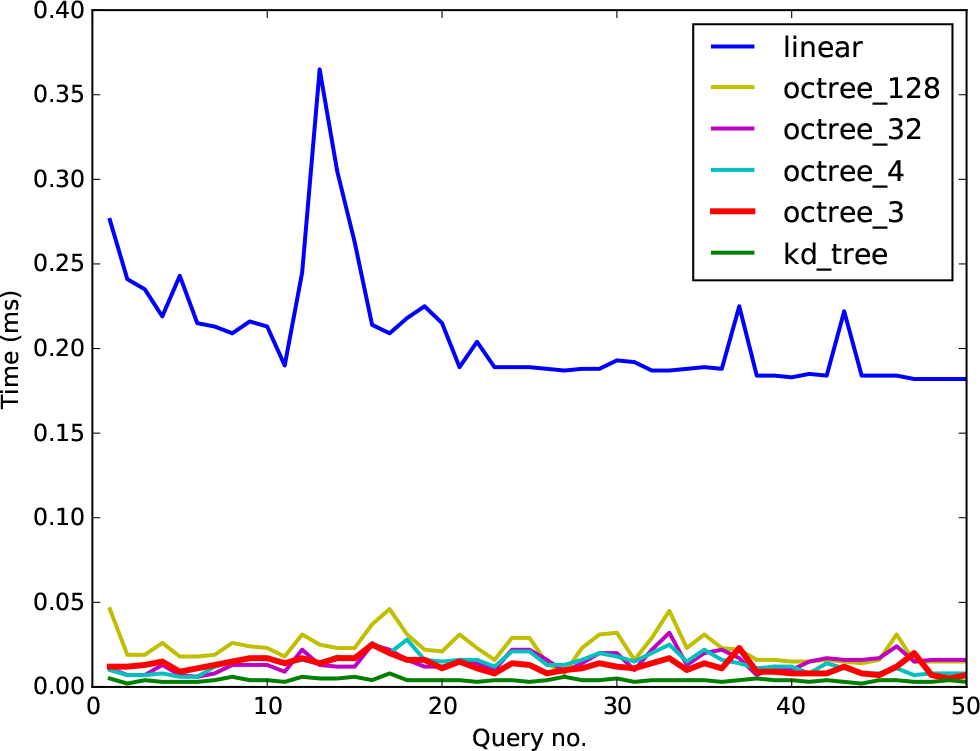
Comparison of kD-Tree and Octree. kD-Tree is most efficient for neighbor searching. Octree is slightly slower, for dynamic or semi dynamic data sets of macromolecules voxel capacity of 3 to 32 is best but it could be improvised based on the nature of data set.

### 3.2. Exception handling

The casual-parse mode in our parser does not check any passing criteria for handling individual attributes of ATOM/HETATM. However, strict-parse mode ensures only those attributes which fulfil the PDB format guidelines for naming and representation of the values. Many faulty .pdb files can be validated using this method at parsing stage. We were able to detect a few such .pdb files using our parser, some of them are also reported to the RCSB database administrators. There are other faults which we have tried to overcome; labeled as PDB format faults, structural faults and uncorrectable faults.

1. We have attempted to correct and overcome some of the PDB format limitations in our program. We have parsed all the important header section data and stored in our data structure which is easily accessible. Using various ligand names for single ligand in PDB file highly discouraged, for example water molecule is found as H2O, WAT, HOH, etc but it is simplified into a single name as HOH.
2. The structural faults are identified and reported; showing un-natural dihedral angles, giving contradictory UNIPROT and structural sequence alignments, removing solvents which are present because of experimental techniques. Graph data structure specially designed to represent covalent bonds of only highest occupancy atoms and reports missing atoms.
3. We are still left with many faults which are uncorrectable due to errors that stem from experimental data, for instance, the electron density might not match with the coordinates. Plenty of PDB format limitations are intrinsic, these have been addressed in newer formats like mmCIF [17] and mmTF.

## 4. Discussion

### 4.1. Extension for biochemical applications

The developed parser can be used in different aspects of macromolecular system analysis such as: sequence and structure comparison, surface atoms detection, active water molecule identification and salt bridge determination etc. In addition to this, the program is also able to detect the active site in a protein and identify the presence of any metal and its interactions.

We have used this parser in our lab for analyzing numerous macromolecular structures, virtual screening and docking results, non-covalent interaction studies, .pdb files from molecular dynamics simulations and other structure based data-set studies. In a case study (see supplementary material), we are able to identify stabilizing and destabilizing salt bridges from two homologous mesophilic and thermophilic *α*-carbonic anhydrase[17]. Studying these types of structural factors like salt bridges, surface residues, and active site water molecules for any macromolecular system is much more simple, fast and error-free.

The protein’s coordinate, bonding; dihedral and other such structural data can be used to study different structural and functional aspects of it. The new parser, which as discussed so far, provides an innovative, faster and necessary platform to prepare the data in structured form for further analysis. Using which we have incorporated various macromolecular analytical tools at one place. The motivation of our work is to provide a integrative tool for in-silico studies that range from getting FASTA sequence to getting active site information.

### 4.2. Applicability and scope

The user interface provides physical abstractions (e.g. atoms, bonds, molecules) of the data that could be easily manipulated by the user.Having active and growing international developer community, ongoing and future developments will improve performance further, introduce transparent parallelization schemes to utilize multi-core and GPU systems efficiently, and interface with high performance data analytics algorithms [18]. We think that this will be a major step in bringing forward C++ language for biology and it being used for open source for its performance and therefore contributing in improving the popularity and current lack of any ongoing projects.

We, strongly recommend that the format of the .pdb file should be reconsidered for making it software-friendly so as to improve the performance of the software as well as to improve the digital readability of the format. A more detailed and strict organization of the attributes is called for. These attributes should be separated by something like space aiding in the distinction of the values which are currently merged in many cases and, therefore, become vague. Now, for solving this problem, we have developed a efficient parser which can arrange all the important data in a well-mannered form as being a fastest program among various others. In addition to the task of parsing of the .pdb, .cif and .mmtf files for working with static structures for molecular analysis; we also wish to add dynamic molecular structural analysis in future.

### 4.3. Availability

The source is available on bitbucket at https://bitbucket.org/gazalk/pdb_parser/. It is based on C++11 and requires Qt >5.6 and MsgPack for C to be pre-installed. MAT is available as a web-service at http:/mat.iitr.ac.in/.

## 5. Supplementary Material

See Supplementary material for grammar rules and grammar table in Tables S1 and S2 for tokenization of each section in the parser. The benchmarks for hydrogen bond finding are shown in Figure 1. Construction and searching of the data structures, kD-Tree and Octree is given in Figure 2 and Figure 3 of the Supplementary data. The case study of *α*-carbonic anhydrase to mutation design in Figure 4 and Figure 5 while showing non-covalent interactions in Table S3.

## 6. Funding

This research was supported by the Department of Biotechnology (DBT), Ministry of Science and Technology, Government of India [Grant number DBT/ 2015/IIT-R/325] to G.K.

## 7. Acknowledgements

The work was partially conducted in the context of the Bioinformatics Resources and Applications Facility (BRAF), C-DAC, Pune, India. Institute Computer Center, IIT-Roorkee and Bioinformatics Facility, Department of Biotechnology, IIT-Roorkee granted the provision of computational facilities and support.

